# Design of a new model yeast consortium for ecological studies of enological fermentation

**DOI:** 10.1101/2024.05.06.592697

**Authors:** Eléonore Pourcelot, Audrey Vigna, Thérèse Marlin, Virginie Galeote, Thibault Nidelet

## Abstract

Wine fermentation involves complex microbial communities of non-*Saccharomyces* yeast species besides the well-known *Saccharomyces cerevisiae*. While extensive research has enhanced our understanding of *S. cerevisiae*, the development of multi-species fermentation starters has led to increased interest in yeast interactions and the role of microbial diversity in winemaking. Consequently, molecular methods have emerged to identify the different species at different stages of the winemaking process. Model microbial communities or consortia, which provide simplified systems resembling natural microbial diversity, offer opportunities to investigate population dynamics and understand the role of community diversity in ecosystem performance.

Here, this work aims to design a yeast consortium reflecting the diversity of wine yeasts and to develop a method for accurately tracking their population dynamics during fermentation. We developed and characterized a six-species consortium, with *S. cerevisiae*, *Hanseniaspora uvarum*, *Starmerella bacillaris*, *Metschnikowia pulcherrima*, *Lachancea thermotolerans* and *Torulaspora delbrueckii*. By tagging each yeast species with distinct fluorescent markers, the study enables real-time monitoring of individual species within the consortium using flow cytometry. We have carried out a complete analysis of this consortium, studying the evolution of populations over time and examining factors such as metabolite production and fermentation kinetics.

In addition, the yeast consortium was used to test the diversity-function relationship as a proof of concept. We sought to determine the impact of the initial evenness on communities’ performances subjected to osmotic stress. To this end, ten randomly designed consortia with varying initial species proportions were followed in enological fermentation with 200 and 280 g/L of initial sugars. The initial proportion of certain species affected the population dynamics and metabolite production however no demonstrable effect of the initial evenness on the response to osmotic stress was shown.

These results demonstrated the usefulness of the presented consortium, which is now available to the scientific community and can contribute to future work trying to decipher multispecies dynamics and the role of yeast diversity in wine fermentation.

## Introduction

Fermented foods and beverages are consumed worldwide, and their production relies on diversified microbial communities that include numerous bacteria and yeasts (Furukawa et al., 2013; Wolfe & Dutton, 2015; Tamang et al., 2016). Wine alcoholic fermentation involves different yeast species from the *Saccharomyces, Hanseniaspora, Pichia, Metschnikowia, Starmerella, Torulaspora, Lachancea*, or *Rhodotorula* genera (Barata et al., 2012; Drumonde-Neves et al., 2021). Research has mainly been focused on optimizing the process with *Saccharomyces cerevisiae*, as it is the main species carrying out the alcoholic fermentation. Thus extensive knowledge has been provided on *S. cerevisiae* growth, and its influence on wine aroma as well as the genetic basis of its suitability for fermentation (Lambrechts & Pretorius, 2000; Sablayrolles, 2009; Peter et al., 2018). However, the development of multispecies starters, with one or two non-*Saccharomyces* strains alongside *S. cerevisiae* (Roudil et al., 2020), has spurred research on interactions between yeast species and on the role of wine microbial diversity in the process (Ivey et al., 2013; Ciani & Comitini, 2015; Bordet et al., 2020).

Consequently, an increasing number of studies on wine microbial ecology, ranging from investigation on the factors influencing yeast and bacterial diversity in grape must, such as climate and vine management (Bokulich et al., 2014; Bagheri et al., 2016; Grangeteau et al., 2017), to the mechanisms of interactions involved between the different species (Bordet et al., 2020). Studies on population dynamics have highlighted mechanisms of competition for nutrient, secretion of toxic compounds or involvement of cellular contact (Holm Hansen et al., 2001; Medina et al., 2012; Kemsawasd et al., 2015). Molecular mechanisms and phenotypic changes caused by interactions have also been studied through transcriptomic analysis or description of physical interactions of cell wall proteins (Brückner et al., 2020; Mencher et al., 2021; Conacher et al., 2022). In addition, interactions and population dynamics of yeasts can be modulated by environmental conditions, such as the different stress factors they encounter during winemaking. Indeed, across the alcoholic fermentation yeasts are exposed to harsh conditions, encompassing low pH, low oxygen availability, or high sugar content (Heard & Fleet, 1988; Bauer & Pretorius, 2000; Varela et al., 2021). The varied environmental conditions and complex microbiota involved in fermentation makes it difficult to draw conclusion.

Synthetic communities that simplified complexe ecosystems, like soil and gut microbiota or food products, allow to disentangle biotic and abiotic factors influencing community dynamics and functionning (Roy et al., 2014; Blasche et al., 2017; Vrancken et al., 2019). Therefore they have been used to explore complex ecological questions. Synthetic communities can be composed of natural isolates that naturally coexist (Venturelli et al., 2018; Kehe et al., 2019; Alekseeva et al., 2021). They can composed of microorganinsm that are engineered to realize complex biotechnological processes (Shong et al., 2012; Minty et al., 2013) or investigate interactions through crossfeeding induced by auxotrophies (Mee et al., 2014; Giri et al., 2021), and can also include genetic modification allowing to identify and follow species in mixes (Dunham, 2007; Kylilis et al., 2018). Model communities from fermented food, especially wine, have recently became a new model to investigate yeast-yeast and yeast-bacteria interactions in multispecies context (Ponomarova & Patil, 2015; Calabrese et al., 2022; Lax & Gore, 2023). In addition, understanding ecological principles in fermented food and beverages may prove useful to help steer the process and obtain predictable results (Louw et al., 2023). Previous research on microbial communities have shown that higher initial community evenness, namely even initial species abundance, can improve stable functionality or resistance to invasion when submitted to environmental stress (Wittebolle et al., 2009; De Roy et al., 2013). For the winemaking yeast community, (Ruiz et al., 2023) recently tested the relationship between richness (number of species) and fermentation process. However, despite works on different inoculation ratio at the beginning of fermentation (Comitini et al., 2011; Morales et al., 2015), the absence of fine monitoring of microbial dynamics during fermentation leaves many shadow areas about the initial evenness impacts alcoholic fermentation for wine.

While traditional numbering methods like differential cultures, qPCR, and metabarcoding are valuable tools for studying microbial communities, they often suffer from limitations in terms of time, labor, and the ability to provide real-time data. Flow cytometry, on the other hand, offers a rapid and accurate method for monitoring population dynamics in multispecies communities, as demonstrated in recent studies (Rigottier-Gois et al., 2003; Kylilis et al., 2018) and model wine yeast pairwise cocultures and consortia (Longin et al., 2017; Conacher et al., 2020).

The present study aimed to develop a novel six-species consortium representative of wine yeast diversity as well as the method to discriminate its different species population. This consortium consisted of one strain of six different species, modified with a specific fluorescent tag, at different initial abundance. We also verified our ability to accurately measure species proportions in complex mixtures with mock communities and finally used this novel model consortium to test the diversity-function relationship as a proof of concept.

## Material and methods

### Strains and media

In this study, six wine yeasts species were used: *Hanseniaspora uvarum, Lachancea thermotolerans, Metschnikowia pulcherrima, Saccharomyces cerevisiae, Starmerella bacillaris*, and *Torulaspora delbrueckii*. The different wild type strains are wild isolates from natural grape must from Southern France and the transformant genotypes are indicated in Table1. Strains were kept at −80°C in yeast peptone dextrose YPD (1% yeast extract, 2% bactopeptone, 2% glucose) supplemented with 20% of glycerol before being streaked on YPD agar and incubated at 28°C. *Escherichia coli* DH5ɑ bacteria (New England Biolabs), used for plasmid constructions, were grown on LB medium (1% tryptone, 0,5% yeast extract, 0,5% NaCl) supplemented with 100 μg/mL of ampicillin antibiotic (Sigma A9817). For solid media preparation, agar was added to the media at 2 %.

### Fluorescent strains construction

#### Plasmid construcion

The following plasmids were constructed by Gibson assembly (Gibson et al., 2009): pFA6a-link-yEmCitrine-NATMX, pFA6a-link-yomCherry-NATMX, pFA6-TDH3.1kb.Td-mCitrine-NATMX, pFA6-Hu1kb-BFP2-KAN. Gibson assembly were done using the NEB Builder HiFi DNA Assembly Master Mix (New England Biolabs) and transformed into *E. coli* DH5ɑ (New England Biolabs) following the manufacturer instructions. pFA6 plasmid backbone, antibiotic resistance cassettes (*KanMX, hphMX*) and fluorescent protein genes (EGFP, mCherry, mCitrine, mTagBFP2) were obtained from plasmids ordered from AddGene (#44900, #44899, #44645, #44903, **Supplementary Table 1**, **(Sheff & Thorn, 2004; Lee et al., 2013)**). Primers used for the amplification of fragments used for the assembly are listed in **Supplementary Table 2**. Where necessary, homologous sequences of approximately 1kb were amplified from the target species (*T. delbruecki* CLIB3069, *H. uvarum* CLIB3221, *S. bacillaris* CLIB3147). All plasmids were checked by enzymatic digestion (New England Biolabs). Plasmid DNA was extracted from 3 mL of overnight *E. coli* LB (with 100 μg/mL of ampicillin) culture with the NucleoSpin Plasmid extraction kit (Macherey Nagel, Düren, Germany) according to manufacturer instructions.

#### Yeast genetic modification

Genes coding for fluorescent proteins were integrated into the genome in fusion of *TDH3* (glyceraldehyde-3-phosphate dehydrogenase) gene (or its orthologue in non-*Saccharomyces* strains). Cassettes used for transformation were amplified from plasmid matrices listed in **Supplementary Table 1** with a high fidelity enzyme, either the KAPA HiFi kit (Cape Town, South Africa) or Phusion High-Fidelity DNA polymerase (Thermo Fisher Scientific, Vilnius, Lithuania), using primers specific to each species, as listed in **Supplementary Table 3**. PCR amplification products were purified with the PCR clean-up Kit (Machery Nagel, Düren, Germany, REF 740609). In addition*, S. cerevisiae* MTF4798 was tagged with a second fluorescent protein, in fusion of *ENO2* (enolase II), another gene involved in glycolysis. *TDH3* has a strong promoter and is expressed continuously during fermentation in *S. cerevisiae*. Similarly, *ENO2* is also expressed throughout *S. cerevisiae* fermentation, even though to a lesser extent (Puig & Pérez-Ortín, 2000; Peng et al., 2015). Yeast transformation was carried out by electroporation, as described by Pourcelot et al. (2023). Transformants were selected on YPD agar supplemented with the appropriate antibiotic: 800 µg/mL of hygromycin B for *S. bacillaris*, 100 µg/mL of nourseothricin for *S. cerevisiae*, *T. delbrueckii*, and *L. thermotolerans*, and 200 µg/mL of geneticin for *H. uvarum* and *S. cerevisiae*. Integration at the locus was verified with two independent PCR (**Supplementary Table 4**) performed on at least 8 clones. To check that transformation did not modify cells behavior, growth of wild type strains and two transformants were compared by microplate assay (**Supplementary Figure 1**).

### Fermentations

#### Synthetic must

Synthetic grape must was prepared according to (Bely et al., 1990) with 425 mg/L of yeast assimilable nitrogen (as a mixture of amino-acids and ammonium), supplemented with 5 mg/L of phytosterols (Beta-sitosterol, Sigma, St-Louis, US, MO) and a final pH of 3.3. Total sugar concentration was either 200 g/L (S200, with 100 g/L of glucose and 100 g/L of fructose), or 280 g/L (S280, with 140 g/L of glucose and 140 g/L of fructose). Synthetic must was pasteurized at 100°C for 15 min before use.

#### Cell inoculation

Cell cultures were prepared with a first propagation in 5 mL YPD for 18 hours at 28°C, and a second propagation of 1 mL in 50 mL of synthetic must S200 for 24 hours at 28°C with agitation. Cell concentration of each pre-culture was measured by flow cytometry before inoculation. Appropriate cell culture volume was used to start fermentation with a final cell concentration of 10^6^ cells/mL. In the case of consortia, the volume of different species cultures were pooled to obtain the desired initial abundance (**Table 2**). Cell suspensions were centrifuged at 4415 g for 5 min and washed in saline solution (9 g/L of sodium chloride, Sigma), and suspended in 5 mL of synthetic must for inoculation (either S200 or S280).

**Table 1:** List of yeast strains used in this study.rajouter colonne parental strain pour y mettre les souches d’origine.

| Species | Strain name | Parental strain | Code | Genotype* | Provider/Reference** |
| --- | --- | --- | --- | --- | --- |
| *S. cerevisiae* | MTF 4798 | MTF1152 |  | *TDH3-mCherry-kanMX* | This study |
| *S. cerevisiae* | CLIB 3930 | MTF 4798 | Sc | *TDH3-mCherry-kanMX ENO2-GFP-natMX4* | This study |
| *H. uvarum* | CLIB 3118 | - |  |  | CIRM Levures |
| *H. uvarum* | CLIB 3929 | CLIB 3118 | Hu | *TDH3-BFP2-kanMX* | This study |
| *L. thermotolerans* | MTF 5257 | - |  |  | SPO lab |
| *L. thermotolerans* | CLIB 3932 | MTF 5257 | Lt | *KLTH0G15730-mCherry-natMX4* | This study |
| *M. pulcherrima* | CLIB 3329 | - | Mp |  | CIRM Levures |
| *S. bacillaris* | CLIB 3147 | - |  |  | CIRM Levures |
| *S. bacillaris* | CLIB 3933 | CLIB 3933 | Sb | *TDH3-eGFP-hphMX6* | This study |
| *T. delbrueckii* | CLIB 3152 | - |  |  | CIRM Levures |
| *T. delbrueckii* | CLIB 3931 | CLIB 3931 | Td | *TDEL0E04750-mCitrine-natMX4* | This study |

\*Due to lack of information as of writing time, *TDH3* was used for non-*Saccharomyces* to indicate the closest homologous orthologues to *S. cerevisiae* S288C *TDH3* gene sequence when no gene code was available.
\*\***CIRM:** Center for Microbial Resources dedicated to Yeast, CIRM-Levures, SPO, Univ Montpellier, INRAE, MontpellierSupAgro 2, place Pierre Viala – 34 060 Montpellier Cedex 02 France **SPO lab:** Sciences Pour l'Oenologie Univ Montpellier, INRAE, MontpellierSupAgro 2, place Pierre Viala - 34 060 Montpellier Cedex 02 France

**Table 2:** Composition (in %) and diversity index of the different consortia tested in this study. Hu: *Hanseniaspora uvarum*, Lt: *Lachancea thermotolerans*, Mp: *Metschnikowia pulcherrima*, Sb: *Starmerella bacillaris*, Sc: *Saccharomyces cerevisiae*, Td: *Torulaspora delbrueckii*.

| Consortium | Sc | Hu | Sb | Lt | Td | Mp | Shannon index |
| --- | --- | --- | --- | --- | --- | --- | --- |
| *6 species* |  |  |  |  |  |  |  |
| Co | 10 | 35 | 20 | 5 | 5 | 25 |  |
| *5 species* |  |  |  |  |  |  |  |
| C01 | 100 | 0 | 0 | 0 | 0 |  | 0.000 |
| C02 | 5 | 75 | 5 | 10 | 5 |  | 0.895 |
| C03 | 5 | 5 | 15 | 5 | 70 |  | 0.983 |
| C04 | 5 | 25 | 60 | 5 | 5 |  | 1.102 |
| C05 | 5 | 5 | 50 | 35 | 5 |  | 1.163 |
| C06 | 5 | 30 | 5 | 10 | 50 |  | 1.237 |
| C07 | 5 | 23.75 | 23.75 | 23.75 | 23.75 |  | 1.515 |
| C08 | 5 | 25 | 50 | 15 | 5 |  | 1.277 |
| C09 | 5 | 5 | 45 | 20 | 25 |  | 1.327 |
| C10 | 5 | 45 | 25 | 10 | 15 |  | 1.370 |
| C11 | 5 | 25 | 30 | 30 | 10 |  | 1.449 |
| C12 | 15 | 40 | 25 | 10 | 10 |  | 1.458 |
| C13 | 0 | 25 | 25 | 25 | 25 |  | 1.386 |

#### Scale of fermentation

Fermentations of the 6-species consortium (Co) and monocultures were carried out in 1 L fermenters to allow a higher degree of precision in consortium caracterisation. Due to the greater number of fermenters needed to test the function-diversity relationship, the 5-species consortia were carried out in 250 mL fermenters. One liter fermentation were run in 1.2 L fermenters (200mL headspace) with automated weighing every 20 min to evaluate CO_2_ weight loss. Fermenters filled with 1 L of synthetic grape must were aerated for 40 min before inoculation. An internally developed software was used to measure in real time the rate of CO_2_ production (g/L/h) and fermentations were stopped when it was lower than < 0.02 g/L/h, or after 312 hours if fermentation had not finished yet. At 0, 4, 6, 12, 24, 48, 72, 168 hours of fermentation and when fermentation stopped or after 312 hours, 6 mL were sampled for population and metabolite analysis.

Two-hundred and fifty mL fermentations were run in 300 mL fermenters with manual weighing twice to thrice a day. Fermenters were aerated for 20 min before inoculation. At the start and end of fermentation, 5 mL were sampled for both population and metabolite analysis. At 21, 45, 120, 168, 210, 290 h, 1.5 mL were sampled for population analysis only.

#### Inoculation and consortia composition

Precultures of each strain were done by inoculating 5 mL of YPD with colony grown on a YPD agar plate and incubated for 18 hours at 28°C. One milliliter of the preculture was then propagated in 50 mL in synthetic must S200 in 125 mL Erlenmeyer flasks and incubated for 24 hours at 24 °C with shaking (250 rpm). Volume of cell culture of each species was calculated after measuring cell concentration of each strain culture by flow cytometry. Consortia were made by mixing the adequate volume of each species in individual 50 mL Falcon tubes for each fermenter. The cell mixes were centrifuged for 5 min at 4415 g and the cell pellet was rinsed in physiological water once (NaCl at 9 g/L) then thoroughly ressuspended in 5 mL of synthetic must (S200 or S280 accordingly). Fermenters were inoculated at a total 10^6^ cell/mL rate with this final cell suspension.

Composition of the different consortia used in this study can be found in **Table 2**. The 6-species consortia (Co) had the following initial proportions of the six species were: 35% *H. uvarum* (3.5·10^5^ cells/mL), 25% *M. pulcherrima* (2.5·10^5^ cells/mL), 20% *S. bacillaris* (2·10^5^ cells/mL), 10% *S. cerevisiae* (1·10^5^ cells/mL), 5% *T. delbrueckii* (0.5·10^5^ cells/mL), 5% *L. thermotolerans* (0.5·10^5^ cells/mL). The consortia used for testing the diversity-functionality relationship included eleven consortia with the five fluorescent species (noted C02 to C12). These consortia had varying initial relative abundance of the five different species (except for *S. cerevisiae* which had an initial abundance of 5% in C02 to C11, and 15 % in C12) encompassing different levels of evenness (**Table 2**). Diversity was evaluated with the Shannon index with the following formula: − ∑^*n*^ *p*_*i*_ln(*p*_*i*_) where *p*_*i*_ is the relative abundance of species *i*, and *n* the number of species. C12 composition is adapted from the 6-species consortium but without *M. pulcherrima*. Two consortia consisting of equal initial abundance for all species, with *S. cerevisiae* (C07) or without (C13), as well as a control monoculture with *S. cerevisiae* alone (noted C01) were also included.

### Cell numbering by flow cytometry

#### Mock communities

Cell number of each species was monitored by flow cytometry with the Attune NxT™ Thermofisher® Flow Cytometer (Life Technologies, Singapore). Each population tagged with one or two fluorescent protein was detected with a specific set of channels described in **Table 3**. To validate the method, accuracy of the numbering of the 6 species was made with mock communities. The mock communities were constructed by incubating single species cultures overnight in synthetic must at 28°C with agitation. Subsequently, one mL of culture was centrifuged at 4415 g for 5 min and the resulting cell pellet was resuspended in 1 mL of PBS (130 mM NaCl, 2.6 mM KCl, 7 mM Na_2_HPO_4_, 1.2 mM KH_2_PO_4_, pH 7.4; Sigma). Single-species cell suspensions at a concentration of 10^6^ cells/mL were prepared in PBS and combined to create mock communities in 96-well microplates (200 µL of final volume, 10^6^ cells/mL). Measures were run after a 3-fold dilution in a new microplate. In total, 30 communities were tested, each included one species with theoretical abundance of either 1, 5, 10, 50 or 90% while the five other species were in equal proportions. Cell numbering was performed both immediately after preparation of the mock communities in the microplate and after 2 hours at room temperature.

**Table 3:** Description of the set of channels (with dichroic filters details) used for the detection of the different populations considered in this study. Voltages were set using fluorescent and wild type cells to ensure a proper signal of both. The fluorescence phenotype of each strain is indicated with + (fluorescent) and – (non-fluorescent) on the corresponding channel. Viability was measured using Propidium Iodide (PI).

| Population | Fluorochrome | Channel (laser and filter wavelengths in nm) | Voltage | Fluorescence |
| --- | --- | --- | --- | --- |
| Dead cells | PI | BL3 (488 - 695-40) | 340 | + |
| Live cells | PI | BL3 (488 - 695-40) | 340 | - |
| *S. cerevisiae* | mCherry + GFP | YL2 (561 - 600DLP - 620/15) | 320 | + |
|  |  | BL1 (488 - 495DLP-530/30) | 260 | + |
| *L. thermotolerans* | mCherry | YL2 (561 - 600DLP-620/15) | 320 | + |
|  |  | BL1 (488 - 495DLP-530/30) | 260 | - |
| *T. delbrueckii* | mCitrine | BL1 (488 - 495DLP-530/30) | 260 | + |
|  |  | VL2 (405 - 495DLP-512/25) | 400 | - |
| *S. bacillaris* | EGFP | BL1 (488 - 495DLP-530/30) | 260 | + |
|  |  | VL2 (405 - 495DLP-512/25) | 400 | + |
| *H. uvarum* | mTagBFP2 | YL2 (561 - 600DLP-620/15) | 320 | - |
|  |  | BL1 (488 - 495DLP-530/30) | 260 | - |
|  |  | VL1 (405 - 417LP-440/50) | 340 | + |
| *M. pulcherrima* |  | YL2 (561 - 600DLP-620/15) | 320 | - |
|  |  | BL1 (488 - 495DLP-530/30) | 260 | - |
|  |  | VL1 (405 - 417LP-440/50) | 340 | - |

#### Population dynamics during fermentation

At the different sampling timepoints, 200 µL of sample were ressuspended in 200 µL of PBS after centrifugation in 96-wells microplate (V-bottom) at 4415g for 5 min. Samples were diluted in PBS before reading so that the cell concentration was approximately 1·10^5^ to 5·10^5^ cells/mL. Total cell count was measured with the FSC and SSC channels (488 nm laser, 488/10 filter). To restore the fluorescence, cells suspended in PBS were kept at room temperature for 2 hours. This incubation step likely enables the maturation of fluorescent proteins which is oxygen-dependent (Hansen et al., 2001; Zimmer, 2002), while oxygen is rapidly depleted during fermentation (Bardi et al., 1999; Moenne et al., 2014). Moreover, we observed a decrease in fluorescence of eGFP under the control of the *ENO2* promoter in *S. cerevisiae* near the end of fermentation, reducing the accuracy of measurement after 144 hours of fermentation (up to 5% of inaccurate events at 144 hours). This was corrected with an incubation step for the consortia and *S. cerevisiae* monocultures in YPD for 1 hour (Breslow et al., 2008) followed with an incubation step of 1h in PBS. These steps did not impair total cell count or viability (**Supplementary Figure 2**). Viability was assessed after staining cells with 1 µg/mL propidium iodide (PI, stored at 4°C protected from light; Calbiochem). Cell number and percentage were obtained from gatings done on the Attune NxT software.

### Metabolites analysis

For the metabolite analysis, 5 mL of samples were centrifuged 5 min at 4415 g at 4°C to get rid of cells. Supernatant aliquots were kept at −18°C before being diluted with 2.5 mM sulfuric acid (H_2_SO_4_, Merck, Darmstadt, Germany) at 1:6, and centrifuged again at 16 000 g for 5 min. Diluted samples were kept at - 18°C until analysis. Extracellular metabolite and sugar concentrations (glucose, fructose, ethanol, glycerol, acetic acid, lactic acid, pyruvic acid, and α-ketoglutaric acid) were determined by high-performance liquid chromatography (HPLC), following the method described in (Deroite et al., 2018). Analyses were run on the HPLC (HPLC 1260 Infinity, Agilent Technologies, Santa Clara, California, USA) using a Rezex ROA ion exclusion column (Phenomenex, Torrance, CA, USA) at 60°C, with a flow rate of 0.6 mL/min of 2.5 mM H_2_SO_4_. Concentrations of acetic acid, lactic acid and α-ketoglutaric acid were measured with a UVmeter at 210 nm and other compounds with a refractive index detector and chromatograms were analyzed on OPEN LAB 2X software. Fermentation samples were analyzed with technical duplicates and data analysis were carried out on the mean of both replicates.

### Data analysis

Mock communities were analyzed by comparing theoretical cell concentration to the observed cell concentration, as measured by cytometry.

All fermentations were performed in biological triplicates, on three different runs.

Fermentations of the six monocultures and the 6-species model consortium were compared based on 10 fermentation parameters. They included three CO_2_ kinetics parameters - latency (time needed to reach 5 g/L of total produced CO_2_), time of fermentation (tF, time when CO_2_ production rate < 0.02 g/L/h), maximum CO_2_ production rate (Vmax) - and six metabolic parameters, namely the yields of ethanol, glycerol, and acetic, α-ketoglutaric, lactic and pyruvic acids. Metabolite yields were calculated with the formula:

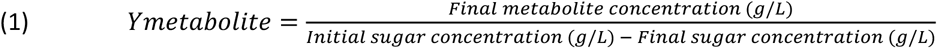

Fermentations of the 5-species consortia were compared with the six metabolites yieds and CO_2_ yield. Differences in yields between the S200 and S280 conditions were visualized by calculating the difference between both conditions with the following formula:

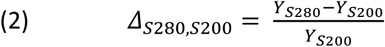

Data were analyzed with R studio software (version 4.3.1, R Core Team, 2024). Mock communities’ data were analyzed with the ‘ggpmisc’ package (Aphalo, 2024). ANOVA were performed to compare the different monocultures and 6-species consortium parameters, followed by a post hoc Tukey test when significant using the ‘rstatix’ package (Kassambara, 2023). For the 5-species consortia, comparison between both sugar conditions were done with t-test for consortia C01 to C13, on the following parameters: residual sugars, yield in CO_2_, and end yields in ethanol, glycerol, and acetic, lactic, α-ketoglutaric and pyruvic acids.

## Results

### Construction of consortium and cytometry numbering optimization

The model community that we used in this study was constituted of 6 different species at different initial abundance: *Saccharomyces cerevisiae* (10% of total population), *Hanseniaspora uvarum* (35%), *Metschnikowia pulcherrima* (25%), *Starmerella bacillaris* (20%), *Lachancea thermotolerans* (5%), and *Torulaspora delbrueckii* (5%). The species and their proportions were determined based on a literature analysis of 18 articles, and the species initial abundance was determined from the average of the initial abundance found in a subselection of 11 articles (Combina et al., 2005; Hierro et al., 2006; Di Maro et al., 2007; Zott et al., 2008; Ocón et al., 2010; Maturano et al., 2015; Ghosh et al., 2015; Bagheri et al., 2017; Simonin et al., 2018; Mateus et al., 2020; González-Alonso et al., 2021). These articles needed to include at least one sampling without inoculation and without enrichment steps, and the identification had to be at the species level. They encompassed different world regions and winemaking practices (grape variety, conventional and organic, etc). Despite the clear influence of these factors on yeast diversity, we found little relationship between the regions and practices with the presence of specific species, which is in line with the conclusions of a meta-analysis (Drumonde-Neves et al., 2021). Among the most frequent species, we decided to select only one species per genera with the assumption that it might increase phenotypic diversity and the overall usefulness of our model consortium. Thus, the six following species were selected: S*. cerevisiae*, *H. uvarum*, *M. pulcherrima*, *S. bacillaris*, *T. delbrueckii* and *L. thermotolerans* (**Supplementary Figure 3A**). Then, the initial abundance for each species was determined from their natural initial abundance in must and early stages of fermentation (**Supplementary Figure 3B**).

The consortium was designed to track individually six species (*H. uvarum*, *L. thermotolerans*, *M. pulcherrima*, *S. bacillaris*, *S. cerevisiae*, *T. delbrueckii*) during fermentation. To this end, our strategy was to label the different species with different fluorescent proteins. The fluorescent proteins were selected based on their excitation and emission spectra to minimize overlaps. Genes of the fluorescent proteins were successfully integrated by homologous recombination in fusion of the *TDH3* gene into the genomes of *H. uvarum* (mTagBFP2), *S. bacillaris* (eGFP), *L. thermotolerans* (mCherry) and *T. delbrueckii* (mCitrine). *S. cerevisiae* was tagged with mCherry fused with *TDH3* and eGFP with *ENO2* (Table 1). For *M. pulcherrima*, a wild type strain was used since we did not manage to integrate genes at the locus even when using long homology sequences. This result is in accordance with previous studies that only obtained random integration and no targeted integration when transforming *M. pulcherrima*, despite the effort of limiting the NHEJ (Non-Homologous End Joining) repair pathway (Gordon et al., 2019; Moreno-Beltrán et al., 2021). As shown on **Figure 1**, which displays derived dot plots and gates used in this study, the different transformants could be well discriminated. *L. thermotolerans* and *S. cerevisiae* are distinguished by the mCherry fluorescence and *S. cerevisiae* by the additional GFP fluorescence. *T. delbrueckii* (mCitrine-tagged) cells could be separated from *S. bacillaris* (eGFP-tagged) using the violet laser, not needing any specific optical configuration, as reported for YFP (Yellow Fluorescent Protein, (Marcus & Raulet, 2013). Initially, the T-Sapphire fluorescent protein was also selected, but its signal overlapped with the mTagBFP2 signal and was not used further.

**Figure 1:**
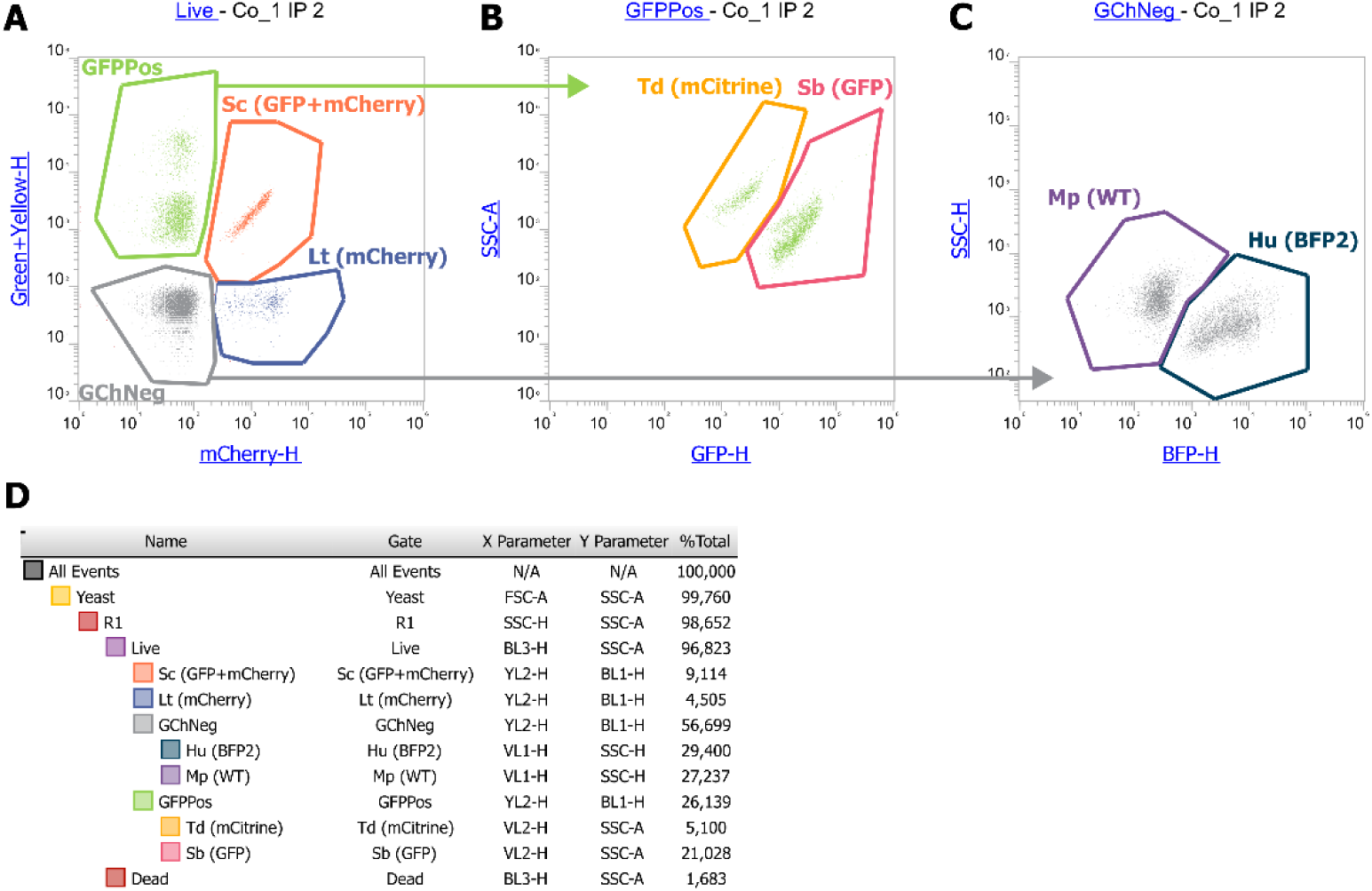
Cytometry set up with an example of sample at inoculation (10^6^ cells/mL at sampling, followed by a 3-fold dilution before the measure). **A:** YL2 (600DLP-620/15)/BL1 (495DLP-530/30) dot plot is used to separate GFP positive and mCherry positive cells. **B:** VL2 (495DLP-512/25)/SSC dot plot derived from the GFPPos gate on A, is used to separate mCitrine (-) from eGFP signal (+). **C:** VL1 (417LP-440/50)/SSC dot plot derived from the GFPNeg gate on A, is used to separate non-fluorescent cells from BFP positive cells. **D:** Hierarchy of gating and percentage of each population. % Total indicates the % of the total population of events in the corresponding gate.

In addition, accurate numbering of the different yeast populations was verified by comparing measured to theoretical cell concentration in mock communities (**Figure 2**). As they are constructed with known composition, mock communities are used to validate methods for microbial quantification, such as metagenomics or cytometry (Tourlousse et al., 2021; van de Velde et al., 2022; Rué et al., 2023). We tested 6-species yeast mock communities composed of one species with theoretical abundance of either 1, 5, 10, 50 or 90%, the five other species in equal proportions composing the rest. Mock communities were constructed from overnight cultures in synthetic grape must (S200) that were diluted in PBS to a 10^6^ cells/mL concentration. In total, 30 communities were tested by mixing the adequate volume of each single-species cell suspension in microplates, with a final cell concentration of 10^6^ cells/mL. The concentration of each species within the mock communities was measured by flow cytometry. When species concentrations were numbered immediately after the mock assembly, the observed and theoretical cell concentrations were well correlated (R^2^ ranging between 0.8 and 0.99, **Figure 2A**). However, during fermentation, a loss of cell fluorescence was observed after 24 hours, especially for the mCherry-tagged cells. To restore fluorescence, samples needed to stay at least 2 hours in PBS at room temperature. This step was also tested on the mock community, and except for *M. pulcherrima* that was under-estimated after the 2 hours (R^2^_Mp_ = 0.73, **Figure 2B**), the other species were still accurately numbered (R^2^ ≥ 0.90).

**Figure 2:**
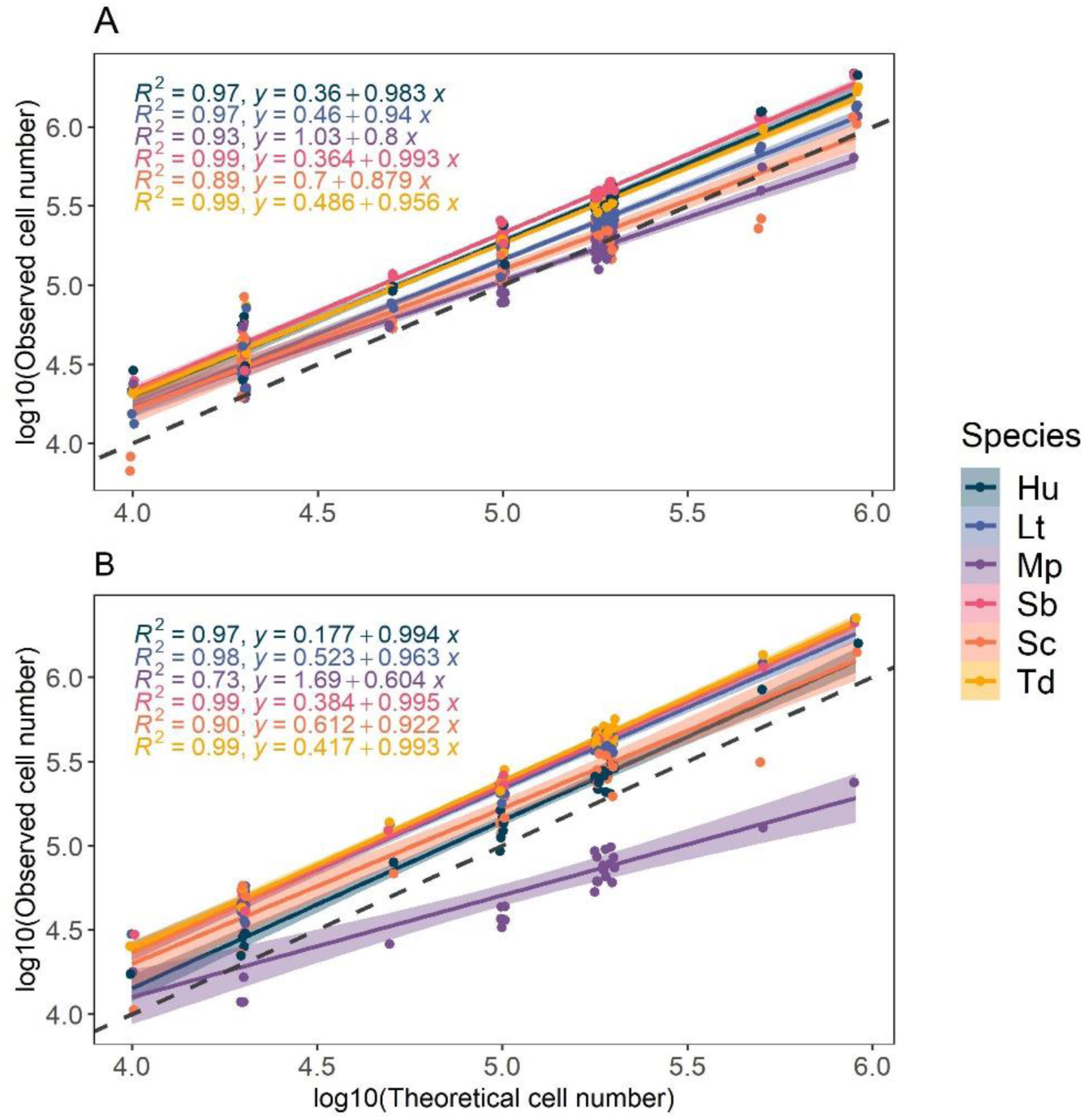
Comparison of observed population and theoretical population in 30 mock communities for immediate measure (**A**) and after 2 hours in PBS (**B**). Correlation factor and linear equation of observed population in function of theoretical populations are indicated for each species. Dashed line: log10(Observed cell number) = log10(Theoretical cell number). Hu: *H. uvarum*, Lt: *L. thermotolerans*, Mp: *M. pulcherrima*, Sb: *S. bacillaris*, Sc: *S. cerevisiae*, Td: *T. delbrueckii*.

### Characterization of the 6-species model consortium

#### Population dynamics in the consortium and monocultures

Monocultures and the 6-species model consortium (Co) were characterized with fermentation in synthetic grape must (S200) at 1 L-scale. All fermenters were inoculated with a total cell concentration of 10^6^ cells/mL and initial abundance of the six species in the 6-species consortium (Co) were 35% *H. uvarum*, 25% *M. pulcherrima*, 20% *S. bacillaris*, 10% *S. cerevisiae*, 5% *T. delbrueckii*, 5% *L. thermotolerans*. The dynamic of species abundance in the consortium are shown in **Figure 3** and population dynamics of monocultures in **Supplementary Figure 4**. In monocultures, most species reached their maximum population between 24 and 48 hours ranging from 1.6·10^7^ to 9.5·10^7^ cells/mL for *M. pulcherrima* and *S. cerevisiae* respectivelly (**Supplementary Figure 4)**. In addition, except for rare events attributed to noise (< 5%), the fluorescent protein expression in genetically modified species was sufficient to identify live cells of each species in their respective gate throughout fermentation. In the consortium, the early fermentation phase was characterized by a rapid growth of *H. uvarum* that reached 65% of the total population at 12 h, with a maximum population of 6.7 log of cells/mL (**Figure 3**, **Supplementary Table 5**). On the contrary, *S. bacillaris* and *M. pulcherrima* abundance dropped under 5% after 12 hours, even though they were inoculated at 20 and 25%, respectively. *L. thermotolerans*, showed a little growth at 24h, reaching 6.3 log of cells/mL, (**Supplementary Table 5**), but quickly decreased after 48 hours. After 24 h, a drop of viability was observed (live cells < 75%) associated with the drop in abundance of non-*Saccharomyces* species, while *S. cerevisiae* became dominant in the consortium and represented 41% of the total population and 58% of the live population (**Figure 3**). This drop in non-*Saccharomyces* viability is unlikely to be directly caused by the ethanol concentration reached at 24h (8.17 ± 1.06 g/L), since it is inferior to the concentrations reached in monocultures (**Supplementary Figure 5**). Maximum population (8 log) in the consortium was reached after 48 h, with almost only *S. cerevisiae* remaining alive (**Figure 3**).

**Figure 3:**
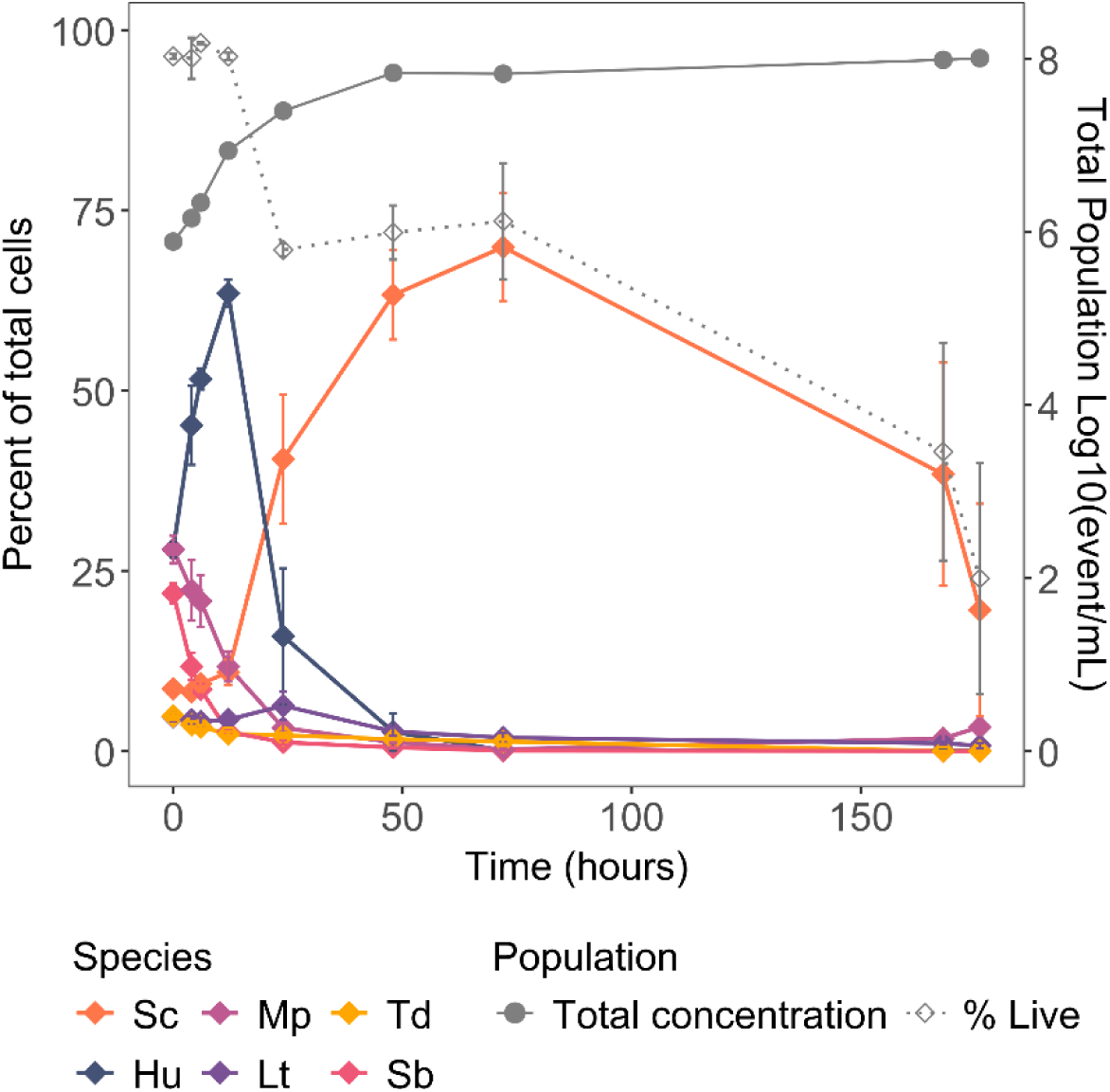
Individual growth of the different species in the 6-species consortium. The grey solid line corresponds to the cell concentration log10(cells/mL), the grey dotted line to the percent of live cell. Solid color lines describe the percent of the different species. Initial abundances were: 10 % *S. cerevisiae* (Sc), 35% *H. uvarum* (Hu), 25% *M. pulcherrima* (Mp), 20% *S. bacillaris* (Sb), 5% *L. thermotolerans* (Lt), and 5% *T. delbrueckii* (Td).

#### Fermentation of the 6-species consortium and monocultures

Fermentation capacity of the 6-species consortium and its member species alone was evaluated through CO_2_ production (**Figure 4**). The consortium performance was similar to that of *S. cerevisiae* alone, with a slightly longer latency and fermentation time due to the lower inoculation rate of *S. cerevisiae* in the consortium compared to the monoculture (latency = 25.3±1.9 h and tF = 158.5±6.1 h compared to 18.7±0.7 and 120.6±3.9 h respectively). The maximum CO_2_ rate (Vmax) of consortium Co was significantly different from any monocultures. It reached 1.74 ± 0.0591 g/L/h (**Figure 4B**), which was between the maximum CO_2_ rate of *S. cerevisiae* (2.16 ± 0.02 g/L/h), *T. delbrueckii* (1.11 ± 0.11 g/L/h) and *L. thermotolerans* (1.01 ± 0.06 g/L/h) alone. The consortium reached the same total CO_2_ production as *S. cerevisiae* (93.08 ± 1.06 g), while *M. pulcherrima*, *H. uvarum*, and *S. bacillaris* alone showed the lowest CO_2_ production (**Figure 4B**).

**Figure 4:**
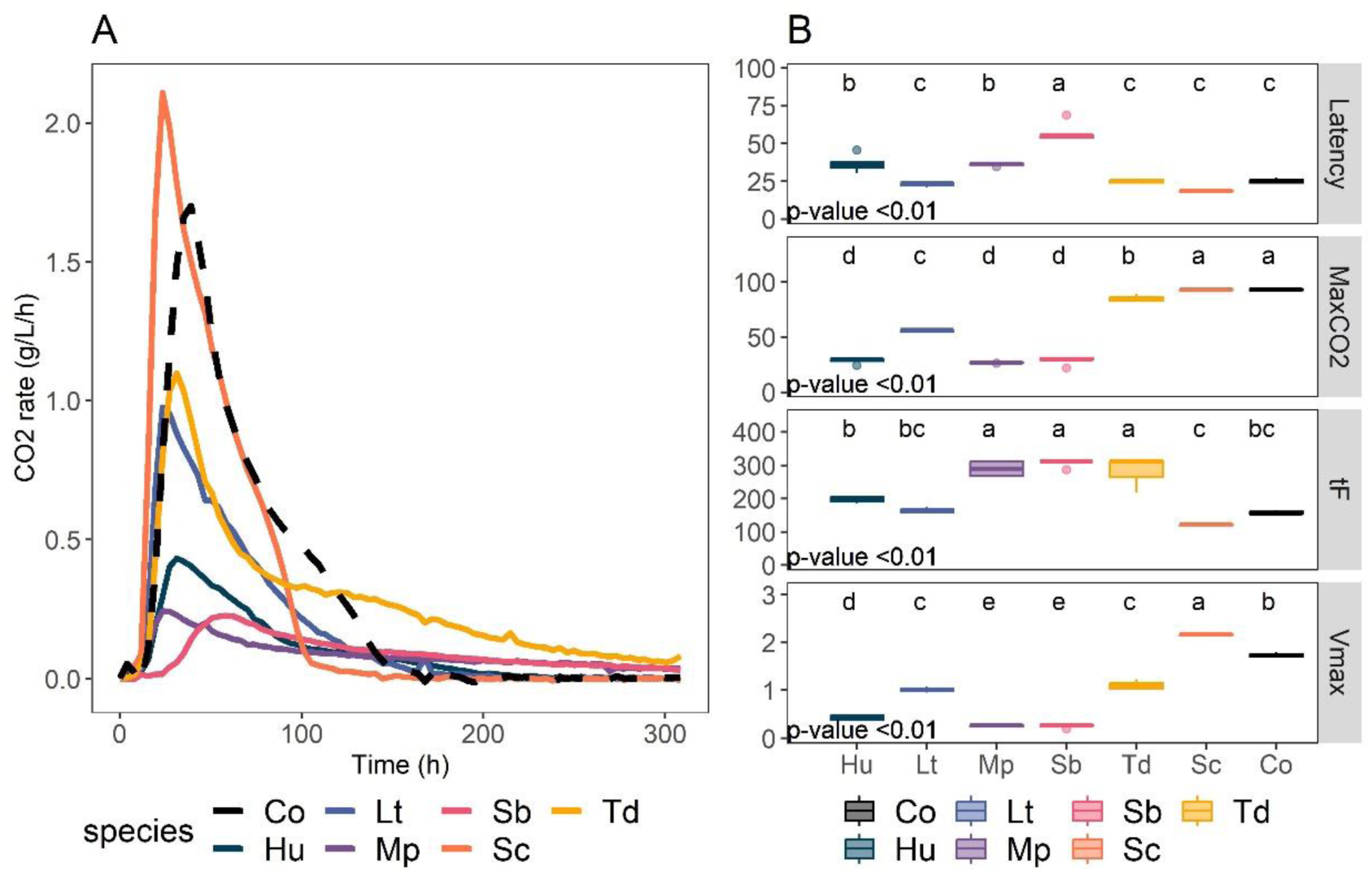
Fermentation kinetics of monocultures and consortium Co. **A:** CO_2_ production rate kinetics of each monoculture (solid line) and consortium Co (dashed line). Smoothing with the Loess method was applied on the results of triplicates. **B:** Kinetics parameters obtained from CO_2_ loss. MaxCO2 = maximum CO_2_ produced (in g/L); tF = fermentation time (time to reach CO_2_ production rate < 0.02 g/L/h), Vmax = maximum rate of CO_2_ production, and Latency = time necessary for total CO_2_ production to 5 g/L. ANOVA results are indicated by the p-value. Statistical groups determined with post-hoc Tukey tests are indicated with lowercase letters. Co: Consortium, Hu: *H. uvarum*, Lt: *L. thermotolerans*, Mp: *M. pulcherrima*, Sb: *S bacillaris*, Sc: *S. cerevisiae*, Td: *T. delbrueckii*.

In addition, changes in concentration of sugars and metabolites from the central carbon metabolism were evaluated. Once again, the 6-species consortium followed the trend of *S. cerevisiae* alone with a delay (**Figure 5**), showing the same end yields in ethanol (0.499 ± 0.001), acetic acid (0.002 ± 0.00), lactic acid (0.003 ± 0.00), and succinic acid (0.003 ± 0.00) as *S. cerevisiae* was alone at the end of fermentation (**Supplementary Figure 6**). Both the consortium and *S. cerevisiae* monoculture reached dryness (total residual sugars < 2g/L), and *T. delbrueckii* left 15.3 ± 5.28 of residual sugars. Other non-*Saccharomyces* left between 80 ± 3.32 g/L for *L. thermotolerans* and 137 g/L of sugars for *M. pulcherrima* and *H. uvarum*. Some difference can also be seen for the α-ketoglutaric acid release by the consortium, which seemed to be produced more slowly than *S. cerevisiae* monoculture (**Figure 5**) but reached a yield similar to monocultures of *S. cerevisiae, L. thermotolerans, T. delbrueckii* or *H. uvarum* (Y_α-keto, Co_ = 1.94e-04 ± 8.66e-06, **Supplementary Figure 6**). *M. pulcherrima* and *S. bacillaris* monocultures overproduced α-ketoglutaric acid in monoculture (Y_α-keto, Mp_ = 28.6e-04 ± 5.9e-04 g/g, Y_α-keto, Sb_ = 15.8e-04 ± 1.4e-04 g/g, **Supplementary Figure 6**). Similarly, *M. pulcherrima* and *S. bacillaris* had the two highest glycerol yields (Y_glycerol, Mp_ = 0.068 ± 0.004 g/g, Y_glycerol, Sb_ = 0.096 ± 0.002 g/g, **Supplementary Figure 6**). Despite that and their high initial abundance in the consortium (45% in total), their early disappearance likely prevented them from producing noticeable amounts of α-ketoglutaric acid and glycerol in the consortium. In the consortium, pyruvic acid was produced in the first 48 hours of fermentation (reaching 0.166 ± 0.005 g/L) then almost entirely consumed at the end (0.023 ± 0.001 g/L, **Supplementary Figure 6**). Monocultures of *S. cerevisiae, L. thermotolerans*, and *H. uvarum* showed a similar behavior with an initial production then consumption of pyruvic acid, reaching respectively a maximum concentration of 0.230 ± 0.013 g/L at 48h, 0.050 ± 0.008 at 24h, 0.153 ± 0.007 at 48h (**Figure 5, Supplementary Figure 6**). On the contrary, *S. bacillaris* and *T. delbrueckii* seemed to produce pyruvic acid without switching to consume it (**Figure 5**). *S. bacillaris* is already known to reroute metabolism towards pyruvic acid production and glycerol for redox equilibrium (Magyar et al., 2014; Englezos et al., 2018). The absence of pyruvic acid consumption may also be related to differences in nutrient requirements, since both nitrogen or vitamins have been reported to modify the production balance of pyruvic acid in *S. cerevisiae* (Fleet, 1993).

**Figure 5:**
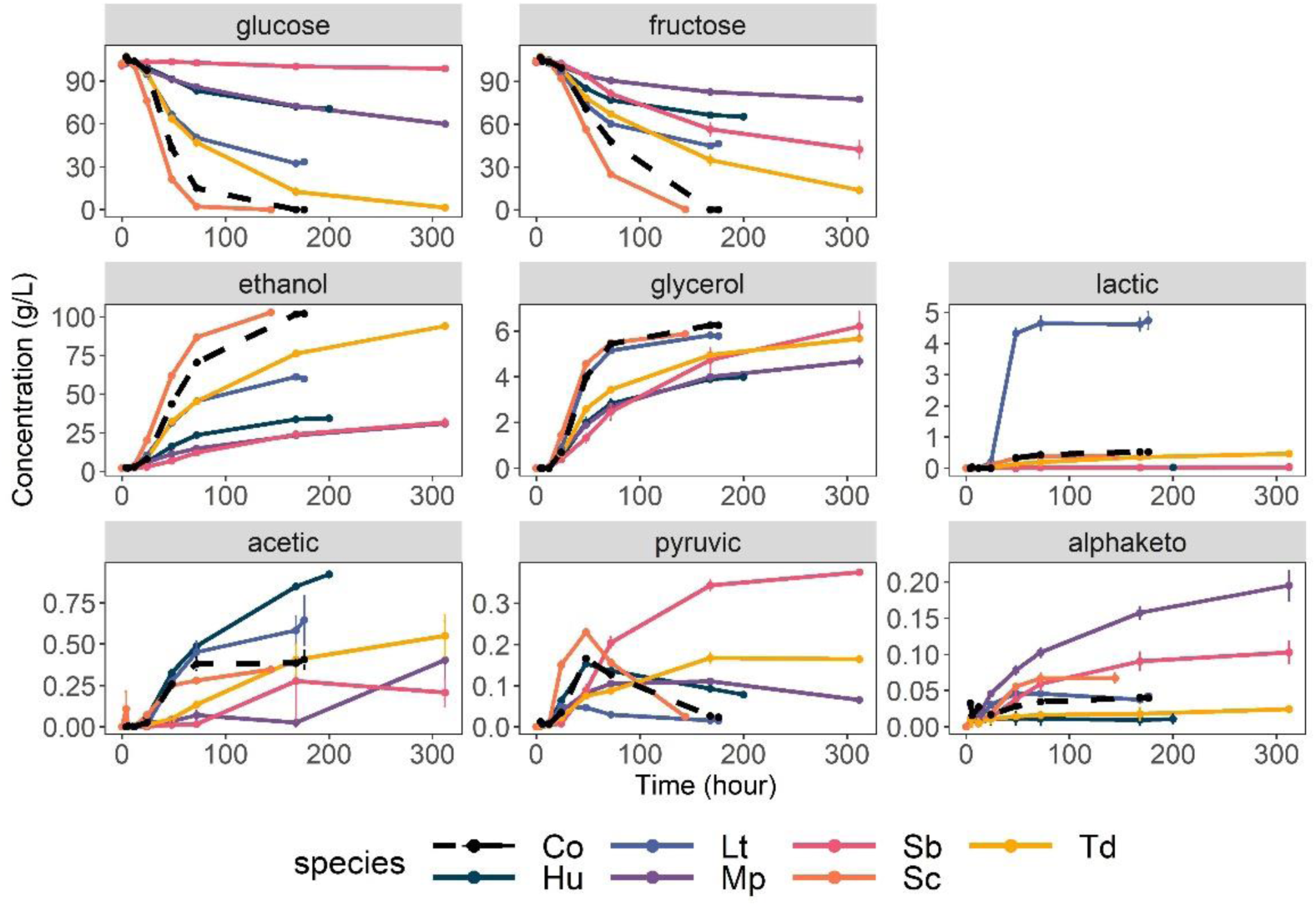
Kinetics of consumption and production of central carbon metabolites in monocultures and consortium. Error bars indicate the standard error. Co: Consortium, Hu: *H. uvarum*, Lt: *L. thermotolerans*, Mp: *M. pulcherrima*, Sb: *S. bacillaris*, Sc: *S. cerevisiae*, Td: *T. delbrueckii*.

### Impact of consortium’s diversity on osmotic stress response

The aforementionned consortium was developped as a tool to further study ecological drivers of the wine microbial community. As a first application, we focused on determining how the initial abundance of species would affect the fermentation process, especially when submitted to an osmotic stress. We tested twelve yeast consortia (numbered C02 to C12) with varying evenness of five species (excluding *M. pulcherrima* due to its underestimation), alongside a *S. cerevisiae* monoculture (C01) and a consortium without *S. cerevisiae* (C13). Fermentations were conducted in 250 mL-scale of synthetic grape must with two sugar concentration: 200 g/L (S200) and 280 g/L (S280). These two concentrations were chosen to simulate a common concentration of sugar found in musts (200 g/L) and a high sugar concentration similar to those already met in some world regions due to climate change (Bock et al., 2013; Gambetta & Kurtural, 2021).

In all consortia initially containing *S. cerevisiae*, this species largely dominated after 120 hours in both sugar conditions and constituted the only live cells, except in consortium C03 (**Figure 6, Supplementary Figure7**, **Supplementary Table 6)**. Unlike the others, in the C03 consortium there was 17% remaining of *T. delbrueckii* at 120h in condition S280, which led to a delayed dominance of *S. cerevisiae* that reached over 95% of total cells only after 168 hours instead of 120 hours in the S200 condition. *S. cerevisiae* monoculture was not much affected by the different sugar conditions and reached its maximum population of 8.12 log at 21 hours (**Supplementary Table 6**). On the other hand, observations revealed variations in the dynamics of non-*Saccharomyces* populations depending on the initial sugar concentration (**Figure 6**) with significant differences in population abundance between S200 and S280 found at 45 hours. In most consortia, *H. uvarum* reached a similar maximum proportion at 21 h for both sugar conditions but decreased more sharply at 45h in condition S280 (**Figure 6**). Similarly, *L. thermotolerans* tended to decrease more in condition S280, except in consortium C13 (without *S. cerevisiae*), while *T. delbrueckii* tended to maintain a higher proportion in condition S280 as compared to S200 (**Figure 6**). At 45 hours in consortium C03, the decrease in abundance between S200 and S280 of *L. thermotolerans* partially matched the increase in abundance of *T. delbrueckii* (6% less *L. thermotolerans* and 15% more *T. delbrueckii* in condition S280). The C03 consortium also showed a longer persistence of *T. delbrueckii* with S280 compared to the other consortia. Interestingly, in the *S. cerevisiae*-free consortium (C13), while both *L. thermotolerans* and *T. delbrueckii* were present at the end of fermentation, *L. thermotolerans* dominated the consortium after 21 hours of fermentation and tended to maintain higher abundance in S280. Regarding *S. bacillaris*, its population decreased very quickly and its proportion was less than 5% after 21h in all consortia, for both sugar concentrations.

**Figure 6:**
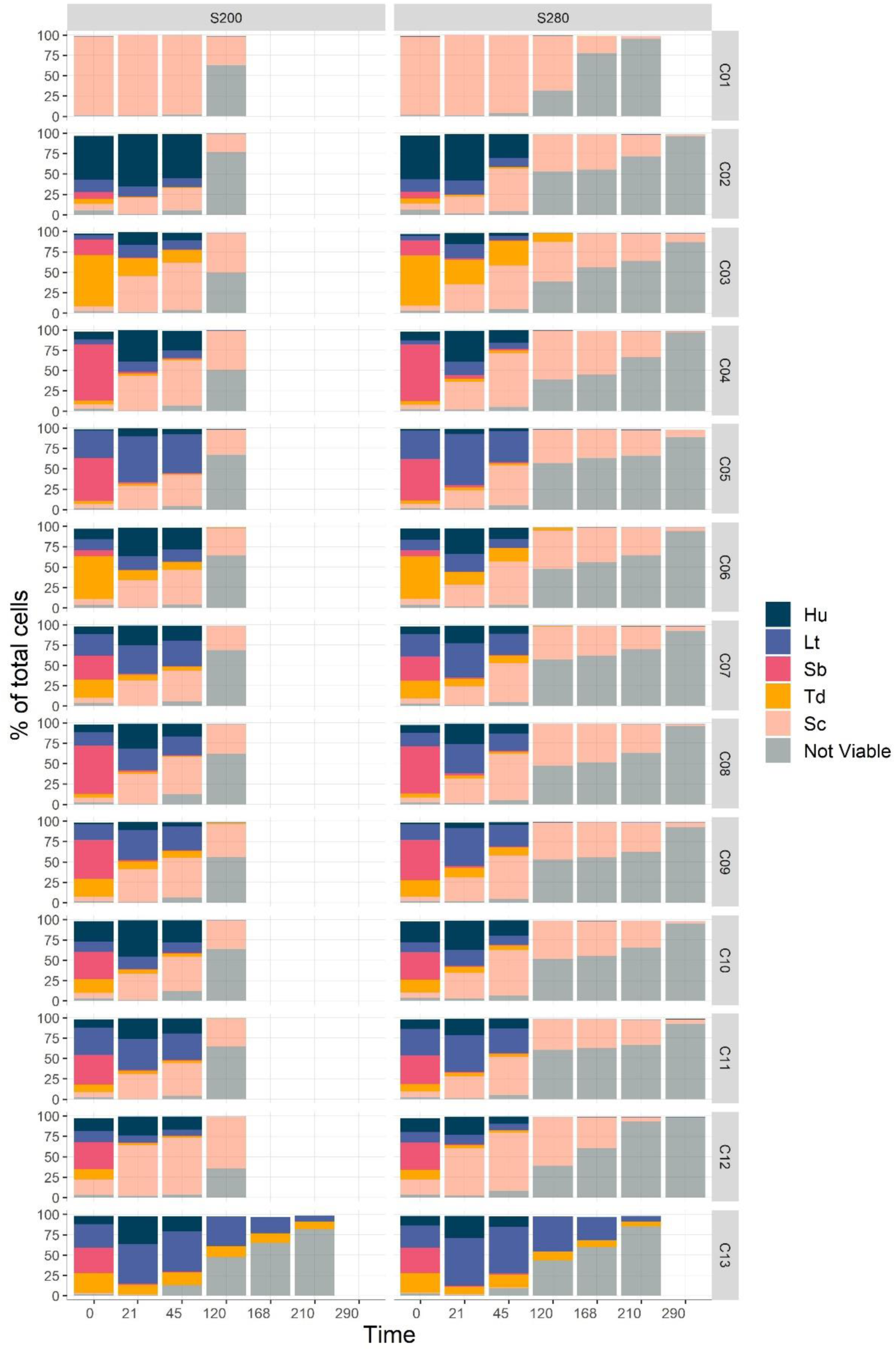
Population dynamics in consortia during fermentation with 200 and 280 g/L of sugars. C01 is a monoculture of *S. cerevisiae*. C02 to C11 initially contained 5% of *S. cerevisiae*. C12 initially contained 15% of *S. cerevisiae* and C13 did not contain *S. cerevisiae*. Hu: *H. uvarum*, Lt: *L. thermotolerans*, Sb: *S. bacillaris*, Td: *T. delbrueckii*, Sc: *S. cerevisiae*.

Fermentation performance was also assessed, through metabolite and CO_2_ yields. The differences in yield of the main metabolite from carbon metabolism between S280 and S200 are shown monocultures in **Figure 7**. Positive values indicate higher yield in condition S280, while negative values indicate lower yield in condition S280.

**Figure 7:**
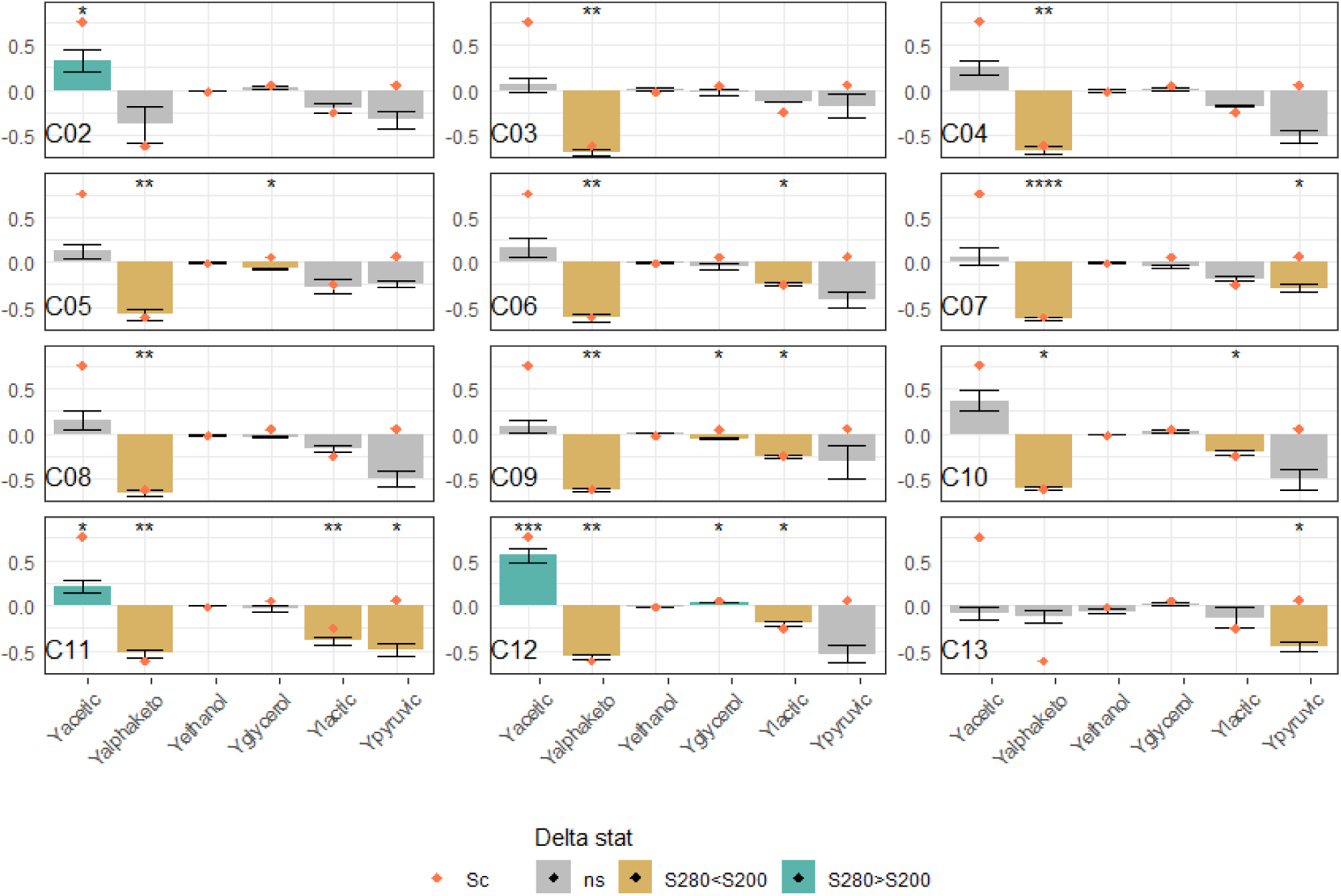
Metabolite yield changes in the different consortia (calculated by dividing the difference between measures for S280 and S200 by the measure for S200). Both sugar conditions were compared by T-test. Gray bars: p-value > 0.05; *, **, ***: p-value inferior to 0.05, 0.01, 0.0001 respectively (in blue when delta is positive, yellow when delta is negative). Orange diamonds indicate the values observed in the *S. cerevisiae* monoculture. C02 to C11 initially contained 5% of *S. cerevisiae*, C12 initially contained 15% of *S. cerevisiae* and C13 did not include *S. cerevisiae*.

Higher initial sugar concentrations led *S. cerevisiae* to produce more acetic acid (increase by 75 % of yield in acetic acid) coupled to a lower production in α-ketoglutaric acid (decrease by 60 % of yield in α-ketoglutaric acid, orange diamond on **Figure 7**).

In addition, we also observed a lower yield in ethanol and lactic acid for *S. cerevisiae* monoculture with high initial sugars, although very small for ethanol (decrease of 1.6 % for the ethanol yield and 25% for the lactic acid yield). We did not observe differences for yield of pyruvic acid for both initial sugar conditions in the *S. cerevisiae* monoculture. We next set out to compare the response to osmotic stress in the consortia. The differences in metabolite yield between the two sugar conditions are shown in **Figure 7**. The different consortia exhibited a wide range of responses, especially regarding the production of acetic and pyruvic acids. Contrary to *S. cerevisiae* monoculture, consortia tended to have lower pyruvic acid yield in condition S280, but differences were significant only in consortia C07, C11 and C13. Regarding acetic acid, its overproduction in condition S280 was greatly reduced in all consortia compared to *S. cerevisiae* monoculture since the highest increase was of 31 % in consortium C02. In the consortium inoculated with a higher proportion of *S. cerevisiae* (C12), 55 % more acetic acid was produced in condition S280 than at S200. In addition, differences in yield of acetic acid in consortia were only significant for consortia C02, C11 and C12. On the contrary, the yield in α-ketoglutaric acid remained lower in condition S280 compared to S200. Differences ranged from 53% lower to 68% lower in consortia C11 and C03 respectively. Only in consortium C02 was it not significantly different between both sugar conditions. This could suggest a prevailing importance of *S. cerevisiae* concerning this metabolite since the difference between both S280 and S200 was often close to that observed in *S. cerevisiae* monocultures, as indicated by the orange diamonds (**Figure 7**). Regarding ethanol, consortia showed an overall lower yield with S200 compared to *S. cerevisiae* monocultures (between 0.482 ± 0.001 in C05 and 0.494 ± 0.003 in C10. Surprisingly, even though glycerol is an essential osmolyte involved in osmotic stress resistance, our results did not show significant differences in glycerol yields between both sugar conditions (**Figure 7**). Our observation may be explained by an unsifficient difference between sugar conditions, and therfore osmotic pressures differences. This too small a difference may not be enough to cause noticeable change in glycerol yield, even though such concentration have been found to cause differences in the lag phase of *S. cerevisiae* alone (Ferreira et al., 2017).

#### Effect of initial composition on consortia behavior

Consortia C02 to C11 were designed to represent variable α diversity, through increasing evenness of the initial relative abundance of non-*Saccharomyces* species (**Table 2**). Hence, the Shannon index of the different consortia ranged from 0.895 in C02 to 1.449 in C11, except C07 that presents an equal proportion of each non-*Saccharomyces* species, and a 1,515 Shannon index. These 10 consortia also included the same initial abundance of *S. cerevisiae* (5%), to avoid changes being related only to *S. cerevisiae*. We were interested in knowing whether an increasing diversity in the yeast consortia would help maintain the fermentation performance during the osmotic stress. However, we found no demonstrable effect of the initial diversity on the different parameters related to fermentation performance, such as yield in CO_2_, yield in metabolites or residual sugars (**Supplementary Figure 8**). However, the initial composition still influenced the behavior of the consortia, especially for the difference in acetic and α-ketoglutaric yields between both sugar conditions (**Supplementary Figure 9**).

## Discussion

In the present study, we constructed a new model community of yeast species that can be numbered by flow cytometry thanks to the expression of different fluorescent proteins. To validate the accuracy of this counting method, we employed mock communities. Then, we characterized the performances of the 6-species community by assessing various fermentation parameters. Finally, to further prove its usefulness, twelve consortia were tested in enological conditions with two differents sugar concentrations to evaluate the influence of varying initial eveness.

In the first part of this work, we tested whether the strategy of species quantification during fermentation was applicable and accurate. Five out of six species were successfully tagged with a set of four fluorescent proteins that can readily be discriminated with a flow cytometry equiped with the appropriate optical configuration. We then assembled 30 mock communities including *S. cerevisiae*, *H. uvarum*, *M. pulcherrima*, *S. bacillaris*, *L. thermotolerans*, and *T. delbrueckii* with abundance ranging from 1 to 90 %. In the range of concentrations tested here, our results indicate that the concentration of cells measured by cytometry highly correlates with the theroretical value. Hence, the quantification strategy could be validated on a consortium including 5 to 6 species. It should still be noted that the accuracy in numbering of *M. pulcherrima* decreased with longer handling time, which could lead to its understimation and should be taken into account when including this species.

During fermentation, species succession of the 6-species community described in this study reflected trends found in both natural communities and laboratory cocultures. For example, as observed in our data, several studies have reported a fast multiplication of *H. uvarum* in early fermentation, including studies in cocultures and consortia (four species) grown in synthetic grape must (Lleixà et al., 2016; Harlé et al., 2020), but also at industrial scale non-inoculated fermentations (Combina et al., 2005). However, in industrial conditions, this increase of *H. uvarum* can be limited when treating must with SO_2_, which is a chemical commonly used as an antimicrobial and antioxidant agent, promoting *S. cerevisiae* (Grangeteau et al., 2017). After 24 hours, *M. pulcherrima* and *S. bacillaris* were detected under 5 % in our consortium. This early decline of *M. pulcherrima* in fermentation has been observed previously (Combina et al., 2005; Chasseriaud et al., 2018), whereas several studies have found *S. bacillaris* able to multiply and maintain a relatively high population in cocultures. For instance, in semi-industrial scale fermentations, *S. bacillaris* could be found in mid fermentation. However, this was related to a later onset of *S. cerevisiae* (Ocón et al., 2010). For other model consortia containing 8 species, tested in both synthetic and natural grape must, *S. bacillaris* remained present up to the mid or end of fermentation (Bagheri et al., 2017). Discrepancies on the presence of *S. bacillaris* during fermentation might be due to different media composition, as was demonstrated with different initial sugar and nitrogen content (Lleixà et al., 2016). In addition, the result of interactions may vary across strains of the same species (Wang et al., 2016; Onetto et al., 2021), even though we observed similar dynamics in cocultures with three different strains of *S. bacillaris* (Pourcelot et al., 2023).

Regarding *L. thermotolerans* and *T. delbrueckii*, both were detectable up to 72 hours in our consortium despite their low inoculation ratio, showing a rather good persistence. In a four species consortium, these two species were able to reach a sizable population despite the presence of *S. cerevisiae.* However, their inoculation ratio was much higher (Conacher et al., 2020). Similarly, *L. thermotolerans* was also found at the end of fermentation of Chenin Blanc grape must, in a consortium of eight species (Bagheri et al., 2017). In natural communities, their maintenance is harder to evaluate since they are present in only low abundance. However, they have been found mid-fermentations (Sternes et al., 2017) and at the end of fermentation in some occasions (Díaz et al., 2013; Simonin et al., 2018).

As for *S. cerevisiae*, it was the main viable species after 24 hours and the only viable species present in the consortium after 72 hours. We can assume that the early drop in viability that we observed between 12 and 24 hours is likely due do the death of non-*Saccharomyces* while *S. cerevisiae* reached its maximum population. Indeed, previous works on cocultures found similar results with non-*Saccharomyces* (such as *L. thermotolerans*, *T. delbrueckii* or *M. pulcherrima)* populations tended to decrease once *S. cerevisiae* reached a high cell number or a high cell number of live *S. cerevisiae* was added (Nissen et al., 2003; Kapsopoulou et al., 2005; Comitini et al., 2011; Taillandier et al., 2014). This phenomenon is also evidenced in sequential inoculation when non-*Saccharomyces* stop growing or even die after the addition of *S. cerevisiae* (Benito et al., 2015; Binati et al., 2020; Seguinot et al., 2020). In addition, a direct toxicity of ethanol can be ruled out as the ethanol concentrations present in the consortium during the non-*Saccharomyces* viability decline were lower than those observed in mono-cultures (**Supplementary Figure 8**) which was also observed by (Holm Hansen et al., 2001). However, recent works from (Lax & Gore, 2023) showed that increasing concentrations of ethanol (even at relatively low concentrations) change bistable interactions into competitive interactions in favor of *S. cerevisiae* highlighting the role of environmental changes on microbial dynamics.

Our data also presented species-related features, such as high glycerol and α-ketoglutaric acid production for *M. pulcherrima* or *S. bacillaris* monocultures. These characteristics have previously been described in monocultures in synthetic must (Englezos et al., 2018; Mbuyane et al., 2022) and in mixed fermentations (Comitini et al., 2011; Englezos et al., 2019; Binati et al., 2020; Seguinot et al., 2020). Similarly, the great production of lactic acid observed here in *L. thermotolerans* monocultures is already known, even though it varies greatly between strains and conditions (Benito et al., 2015; Vaquero et al., 2020). Concerning the 6-species community, its metabolite production and fermentation kinetics seemed mostly influenced by *S. cerevisiae* but was intermediate with non-*Saccharomyces*, which is commonly observed in cocultures (Renault et al., 2013; Bagheri et al., 2017; Harlé et al., 2020). Our second experiment also highlighted the influence of the different species on the fermentation performance in consortia leading to variation in the response to osmotic stress more or less influenced by the presence of *S. cerevisiae*.

As a proof-of-concept, we employed the yeast community with the 5 fluorescently tagged species to investigate the effect of different initial eveness of consortia on their response to two sugar concentrations (S200 and S280). These two concentrations were chosen to simulate a common concentration of sugar found in musts (200 g/L) and a high sugar concentration (280 g/L, (Ferreira et al., 2017; Schmidt et al., 2020). The high sugar concentrations have already been found in different world regions in recent years (Bock et al., 2013; Gambetta & Kurtural, 2021). Altogether, our results indicated a tendency of *T. delbrueckii* and *S. cerevisiae* to have high competitive advantage during fermentations with high sugar content, while *H. uvarum* and *L. thermotolerans* were negatively impacted. *T. delbrueckii* is described as relatively osmotolerant, to both salts and reducing sugars (Lages et al., 1999; Hernandez-Lopez et al., 2003) and was shown to maintain a high population in natural grape musts with 250 to 360 g/L of sugars (Bely et al., 2008; Belda et al., 2015). Thus, the fact that it tended to have a higher competitive advantage at S280 in our study was to be expected. On the other hand, we found that *S. bacillaris* abundance decreased very quickly even in the S280 condition. This is somewhat surprising given this species is osmotolerant and could be expected to have a competitive advantage in high sugar must (Tofalo et al., 2012; Csoma et al., 2020). *S. bacillaris* is also often isolated in fermentations of high sugar must, such as icewine or botrytised wine that can usually contain at least 350 g/L of reducing sugar (Magyar & Soós, 2016). It can even dominate early and middle stages of such fermentations (Nisiotou et al., 2007; Magyar & Soós, 2016). Our inconsistent results might be due to the use of synthetic must with high nitrogen content that could limit the establishment of *S. bacillaris* in a community, in favor of faster growing species. Fewer data are available for *H. uvarum* or *L. thermotolerans* in conditions of high-sugar fermentation. In botrytised must, *H. uvarum* was shown to decrease more quickly than in healthy must, but other factors than the initial osmotic stress, such as presence of other species, may also explain these results since this observation stayed true even when there was no difference in initial sugar (Nisiotou et al., 2007; Contreras et al., 2015).

In the osmotic stress condition, the most striking difference between consortia and *S. cerevisiae* monocultures was the overproduction of acetic acid. *S. cerevisiae* overproduced acetic acid in the S280 condition, while the acetic acid overproduction was smaller in consortia and varied significantly between consortia. This may be partly due to the presence of non-*Saccharomyces* since several studies have reported a positive effect of mixed fermentation of *S. cerevisiae* with non-*Saccharomyces,* namely *S. bacillaris* and *M. pulcherrima,* in reducing the acetate yield (Rantsiou et al., 2012; Contreras et al., 2015; González-Royo et al., 2015; Mbuyane et al., 2022). Another explanation might also lie in the population changes between both sugar conditions. With S280, *H. uvarum* decreased more quickly while *T. delbrueckii* stayed longer. *T. delbrueckii* is known to produce low amount of acetic acid (Bely et al., 2008; Renault et al., 2009), while *H. uvarum* is associated with high production in acetic acid (Ciani & Maccarelli, 1998; Capece et al., 2005; Mendoza et al., 2019). Their difference in dynamics between both sugar concentrations may therefore influence the overall response.

Our data also showed differences of population dynamics with varying initial composition, which highlights that initial abundance is determinant in the species succession in yeast communities (Lax & Gore, 2023; Conacher et al., 2024). However, we found no correlation between initial evenness and fermentation parameters. Several factors could explain this result. The enhanced functionnality observed when increasing diversity relies on the assumption that more diverse community will include complementary species using various resources through increased richness (Cardinale, 2011). In our case, as we focused only on initial evenness with a constant number of species (and therefore species richness), the range of diversity tested might have been too limited. For example, Ruiz et al. (2023) showed a negative effect of an increasing richness in wine yeast communities. They observed that *S. cerevisiae* could not finish fermentation in some communities due to the higher probability of the presence of antagonistic species. Diversity effect has also been observed in microcosms, where fermentation performance was reduced in the highest and lowest diversity context, due to a lower establishment of *S. cerevisiae* (Boynton & Greig, 2016). In addition, the use of synthetic media may not be sufficient to generate a set of ecological sub-niches that are all the more covered as microbial diversity increases (Hooper et al., 2005; Cardinale, 2011; Shade et al., 2012) and account for multiple stress factors. This seems to lead to discrepancies of results between synthetic media and more realistic substrate (Barny et al., 2024). In an enological context, Schmidt et al. (2020) observed differences in *S. cerevisiae* strains dynamics in natural must with 200 or 280 g/L of sugars, but no difference in synthetic must. The low number of carbon sources present in high concentration might also have led to an increased competition between the species leading to the dominance of a single species (Ratzke et al., 2020). With *S. cerevisiae* having a particulartly high competitive advantage in fermentative environments and being able to actively modify its environment (Goddard, 2008; Williams et al., 2015), the overall fermentation process in consortia was probably driven by *S. cerevisiae* since in both sugar conditions non-*Saccharomyces* species had been outcompeted about 48 hours. The quick decline of non-*Saccharomyces* could limit their impact on the fermentation and thus the actual impact of their initial evenness.

Taken together, our results confirm the applicability of the combined use of fluorescently tagged cells and cytometry to follow several species individually in œnological fermentations. As highlighted by previous papers, working with strains tagged with fluorescent proteins enables real-time tracking of the different subpopulations and has been successfully applied in mixed fermentations of one non-*Saccharomyces* and *S. cerevisiae* (Petitgonnet et al., 2019; Harlé et al., 2020), but rarely included more species (Conacher et al., 2020). The increased number of species that can be included in the community compared to previous fluorescent-strains based model could enable testing more complex community composition. Other studies including more species in their consortium employ cultivation or molecular biology methods to follow the microbial dynamics (Lleixà et al., 2016; Bagheri et al., 2017; Chasseriaud et al., 2018). These methods allow enumeration in natural community without specific tagging of the different species but involve heavier experiments and a longer time between sampling and results. To scale-up the number of community tested across a wider range of environmental conditions, our community should be compatible with microfluidic tools based on fluorescent microscopy (Kehe et al., 2019). In its current form, our approach has three important limitations. First, this approach needs a significant initial time investment in molecular biology techniques to construct the strains and some species still prove difficult to transform precisely. Second, our system is more accurate for sub-populations with high cell density. Indeed, despite our effort, we observed noise events leading to detection of cell from species not present up to 5·10^5^ cells/mL. This limits the current system to high cell density numeration and relative abundance. Third, it is necessary to have a cytometer with the appropritate lasers to detect all the fluorescent proteins used here, which can be limiting for some labs. This might be circumvented by changing the tagging strategy and exploiting various level of expression of only two fluorescent proteins (Anzalone et al., 2021). However, this has never been tested in non-*Saccharomyces* species.

## Conclusion

In this study, we successfully developed a 6-species consortium of fluorescently tagged wine yeasts that allow accurate real-time tracking of species subpopulations during fermentation. The consortium was also fully characterized in terms of population dynamics and phenotype, such as metabolite production and fermentation kinetics. The strains constructed in this work are available to the scientific community at the CIRM-levures. This model community not only includes common species among the most frequent found in grape musts but also species known for their various applications, such as starter cultures, bioprotection, and other biotechnological processes and for which molecular tools are still sparse. This microbial consortium could be a valuable resource for applications in yeast ecology and support future research aimed at investigating the influence of stress factors relevant to the fermentation process, particularly when using multispecies starters. Consequently, the trends observed within the 6- and 5-species communities and our methodology could be applied and tested under a wider spectrum of environmental conditions. Model communities could also include different yeast strains or bacterial species to follow both the alcoholic and malolactic fermentations. Ultimately, the insights gained from studying population dynamics might provide strategies to mitigate fermentations by managing the yeast population.

## Acknowledgements

We are thankful to the CIRM Levures and C. Neuvéglise for providing the yeast strains, F. Macna for carrying out the HPLC analysis, D. Segond for his support with cytometry analysis.

## Data, scripts, code, and supplementary information availability

All data, scripts and supplementary information are available online in the following zenodo repository: https://doi.org/10.5281/zenodo.14007596; Pourcelot et al., 2024.

